# CryoLigATE: enhancing the resolvability of cryo-EM maps in protein-ligand complexes using deep learning

**DOI:** 10.64898/2026.08.04.742718

**Authors:** Nandan Haloi, Rebecca J. Howard, Erik Lindahl

**Affiliations:** Department of Biochemistry and Biophysics, Science for Life Laboratory, Stockholm University, Stockholm, Sweden; Department of Applied Physics, Science for Life Laboratory, KTH Royal Institute of Technology, Stockholm, Sweden; Department of Physics, Chemistry and Biology, Linköping University, Linköping, Sweden; Department of Chemistry, University of Illinois at Urbana-Champaign, Urbana, IL, United States

## Abstract

Cryo-electron microscopy (cryo-EM) has become a central tool for structure-based drug discovery, yet ligand-binding sites often remain substantially less well resolved than the surrounding protein, limiting reliable atomic interpretation. Although deep-learning methods have substantially improved overall cryo-EM map quality, their predominantly protein-focused training limits their ability to recover ligand density. Here we present CryoLigATE, a deep learning framework specifically designed to enhance densities associated with protein-bound ligands in cryo-EM maps. We curated a chemically and structurally diverse dataset of more than 6,000 protein-ligand complexes from the EMDB and PDB, encompassing drug-like molecules, lipids, steroids, carbohydrates and other ligand classes, and trained a hybrid convolutional-transformer network to enhance local density around binding pockets. During inference, CryoLigATE automatically extracts the target region from a preliminary atomic model, requiring no manual map preparation and completing localized refinement in seconds on a desktop GPU. Evaluation on an independent test set of 649 complexes demonstrates substantial improvements in ligand resolvability, particularly for maps with poorly resolved binding sites, while preserving high-quality experimental densities. The enhanced maps recover chemically meaningful features, including ligand functional groups and topological continuity, enabling more confident atomic modeling. By learning the structural diversity of ligand features, CryoLigATE addresses a longstanding limitation of cryo-EM map enhancement and provides a useful framework for improving structural interpretation and structure-guided drug discovery.

## Introduction

Protein-ligand interactions are essential for understanding many biological processes and pharmaceutical mechanisms. In particular, visualizing these binding modes at the atomic level, including the interactions, conformations, and geometries of both the ligand and protein, can facilitate the design of pharmaceuticals that mimic or refine the chemical structures, interactions, and functions of known binders. Recent advances in single-particle cryogenic electron microscopy (cryo-EM) have enabled the determination of historically challenging protein structures, for example involving lipid-membrane interactions or low conformational stability [1–3]. Still, local map regions corresponding to bound ligands often exhibit lower resolution than the surrounding protein scaffold, creating a bottleneck for structure-based drug design. For instance, it was recently demonstrated in a cryo-EM structure of *β*-galactosidase that the global protein density reached 1.5 Å resolution, but the local ligand density only resolved to 3.0–3.5 Å [4].

Although automated model-building algorithms for protein-ligand complexes have seen considerable success in recent years [5–9], these tools are inherently constrained by the quality of the input map. Therefore, there remains a critical need for computational methodologies that address the fundamental physical challenge of refining and enhancing the underlying 3D density itself prior to model building. Recent AI-based post-processing methods, such as EMReady and DeepEMhancer, have shown remarkable results in improving map resolvability [10–12]. These methods that are predominately trained on protein maps have shown their success of learning protein representations including helix, loop and sheets. However, they are less effective when applied to ligands and lipids that contain a broader and more irregular chemical space. As highlighted by Berkeley *et al.* [13], treating maps with these existing AI methods often deteriorates the model-to-map Q-score at the binding site.

To address this gap, we present cryo-EM ligand AI-trained enhancement (CryoLigATE), a deep learning method specifically engineered to enhance the resolvability and contrast of ligand densities within cryo-EM volumes (Fig. 1). By curating a structurally diverse training dataset of protein–ligand complexes, we ensure the network effectively learns the intricate geometries and chemical environments characteristic of small molecules. During inference, CryoLigATE requires only a localized sub-volume of the experimental map. This extraction is performed automatically by the pipeline, utilizing the coordinates of a preliminary atomic model as a spatial reference to isolate the density surrounding the target binding site. The utility of this approach is exemplified by a representative case from our independent test set: the short prokaryotic Argonaute protein MapSPARTA, which defends cells against pathogens, bound to nicotinamide adenine dinucleotide [14]) (Fig. 1). CryoLigATE outputs a sharpened density map of the binding pocket, enhancing high-frequency structural features for the ligand that were previously ill-defined in the experimental map.

**Fig. 1:**
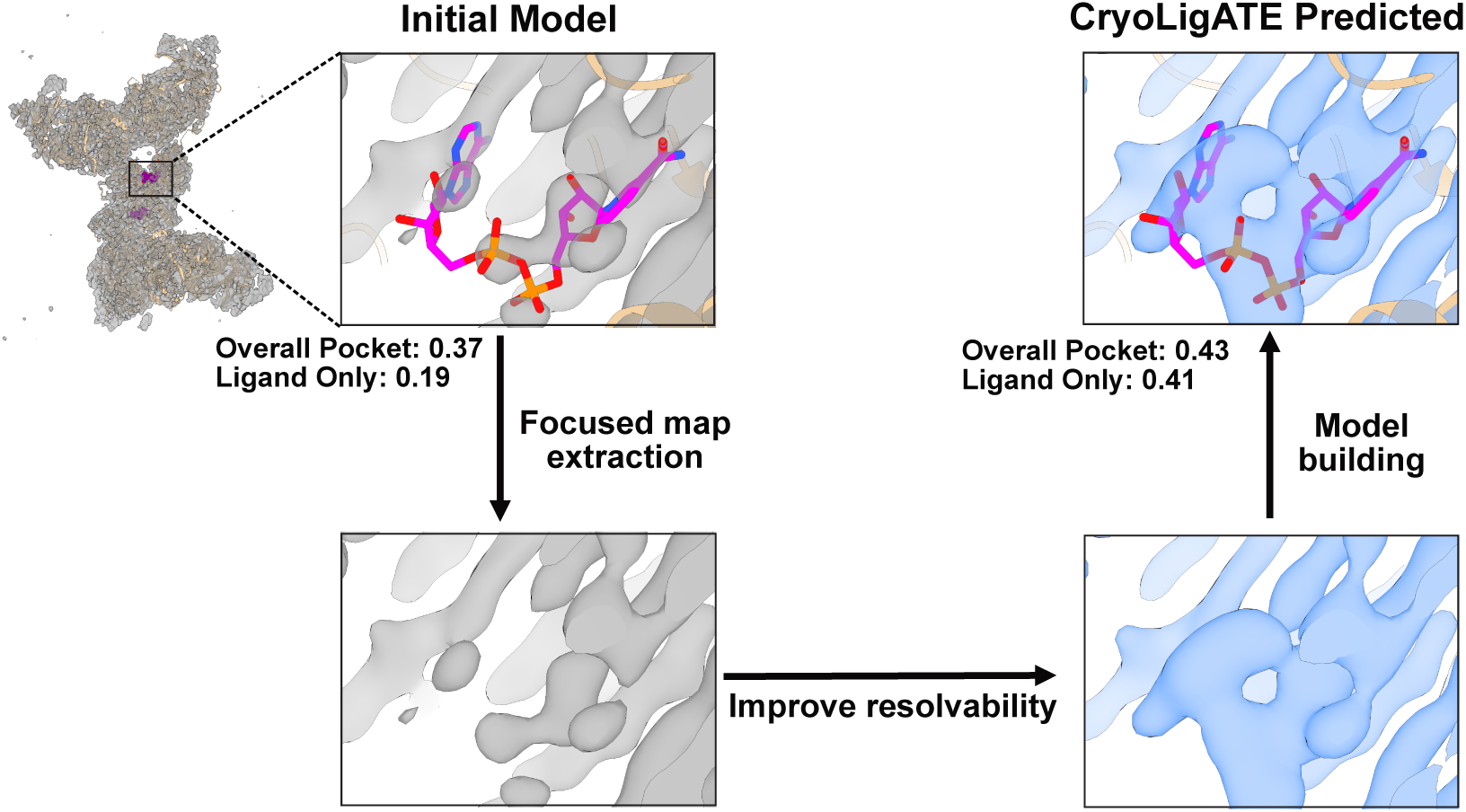
Overview of the CryoLigATE inference workflow. First, a localized 3D density region encompassing the putative binding pocket is extracted from the global consensus cryo-EM map. This extraction is carried out automatically by the pipeline, using the coordinates of an initial atomic model as a spatial reference to isolate the density around the target binding site. This cropped map is provided as input to the CryoLigATE AI model. The model enhances the resolvability and contrast of the local density. The resulting high-resolution map resolves previous ambiguities, facilitating accurate and confident atomic model building for both the ligand and the surrounding protein environment. The entire pipeline can be run with a single inference command line, as described in the Method section. An example of the short prokaryotic Argonaute protein MapSPARTA bound to nicotinamide adenine dinucleotide (NAD^+^) (EMD-40680, PDB 8SPO [14]) is shown from our testing dataset, processed by CryoLigATE.

## Results

### A diverse cryo-EM protein-ligand complex dataset

**Fig. 2:**
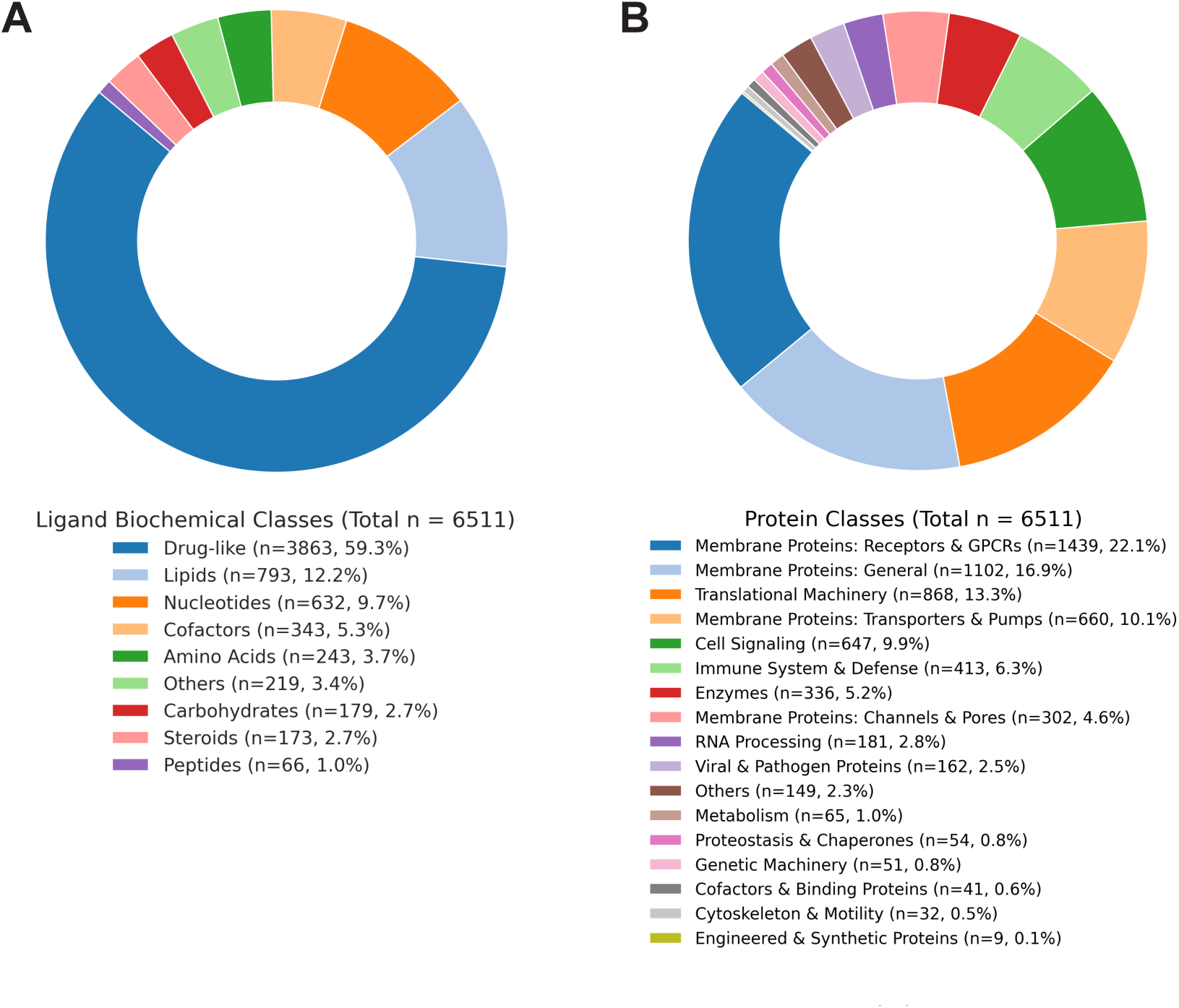
A diverse cryo-EM protein-ligand complex dataset. (A) Distribution of ligand classes determined via an integrated classification pipeline. Drugs were identified by matching InChIKeys against the ChEMBL database (Max Phase *>* 0). Remaining compounds were categorized using RDKit based on SMARTS patterns and physicochemical properties: Lipids (aliphatic chains or logP *>* 4.0), Peptides (*≥* 3 amide bonds), and Drug-like molecules (QED *≥* 0.5 or Lipinski Rule of 5 compliance). (B) Distribution of protein functional families. Classification was performed using a hierarchical keyword-based search of metadata retrieved via UniProt and RCSB PDB APIs.

The curated dataset for training, validating, and testing of our model comprises 6511 unique protein-ligand pairs from cryo-EM structures deposited in the protein data bank (PDB), with an overall resolution of at least 4 Å. It contains diversity across both ligand and protein dimensions (Fig. 2A,B). The ligand population features a prevalence of pharmacologically relevant molecules, with drug-like molecules (59.3%), lipids (12.2%), and nucleotides (9.7%) forming the majority of the set (Fig. 2A). We further analyzed the distribution of physicochemical properties, revealing a diverse structural and chemical space: rotatable bonds range from 0 to 110, ring systems from 0 to 13, molecular weights from 28 to 2975 Da, LogP from -8.86 to 23.3, and Quantitative Estimate of Druglikeness (QED) scores from 0 to 0.95 (Fig. S1).

The protein component of our dataset is also varied, particularly sampling membrane proteins (22.1% receptors, 10.1% transporters, and 4.6% ion channels), various types of translational machinery (13.3%), and enzymes (5.2%) (Fig. 2B). This diversity is reflected in the structural and chemical properties of the binding pockets. Secondary structure analysis of the pockets indicates compositions of alpha-helices (44.8%), beta-sheets (10.4%), random coils (23.0%), and turns (21.7%) (Fig. S2). Furthermore, residues within a 4 Å radius of a given ligand consist of hydrophobic (54.9%), polar (20.4%), basic (10.0%), and acidic (4.6%) amino acids (Fig. S3), underscoring the structural, functional, and chemical divergence of our curated dataset. This dataset is publicly available as described in the Data Availability section for further usage.

### Overview of CryoLigATE

CryoLigATE takes a raw cryo-EM density map as input, and focuses on a specified preliminary ligand, to predict a better resolved map of the protein-ligand complex. For ground truth during training, we used forward-simulated 3D EM maps derived from deposited atomic models as the “high-resolution” targets (Fig. 3A). The EM maps were resampled to a uniform magnification and extracted as 64 x 64 x 64 voxel volumes, corresponding to 32 Å x 32Å x 32 Å. The model employs a 3D Swin-Conv UNet architecture [15] which combines the advantages of conventional residual convolution for local modeling, swin (shifted window) transformer for non-local modeling, and multiscale UNet for further enhancement of local and non-local modeling (Fig. 3B). The transformer combines self-attention of non-overlapping local windows and non-local cross-window connection by shifted window partitioning.

The dataset was partitioned using a random 80:10:10 split for training, validation, and testing, respectively. To eliminate data leakage and ensure a rigorous evaluation, the dataset was partitioned using a sequence-identity-based clustering workflow (see Methods for more details). This resulted in a total of 5195, 667, and 649 data points for the training, validation and testing dataset, respectively. During training, the loss function was optimized by comparing the network’s predicted sharpened density against a simulated target map derived from the atomic model (details in the Method section, Fig. 3). The network was trained for 200 epochs, exhibiting a consistent reduction in both training and validation loss (Fig. S4). The results presented hereafter are derived exclusively from the independent test set.

**Fig. 3:**
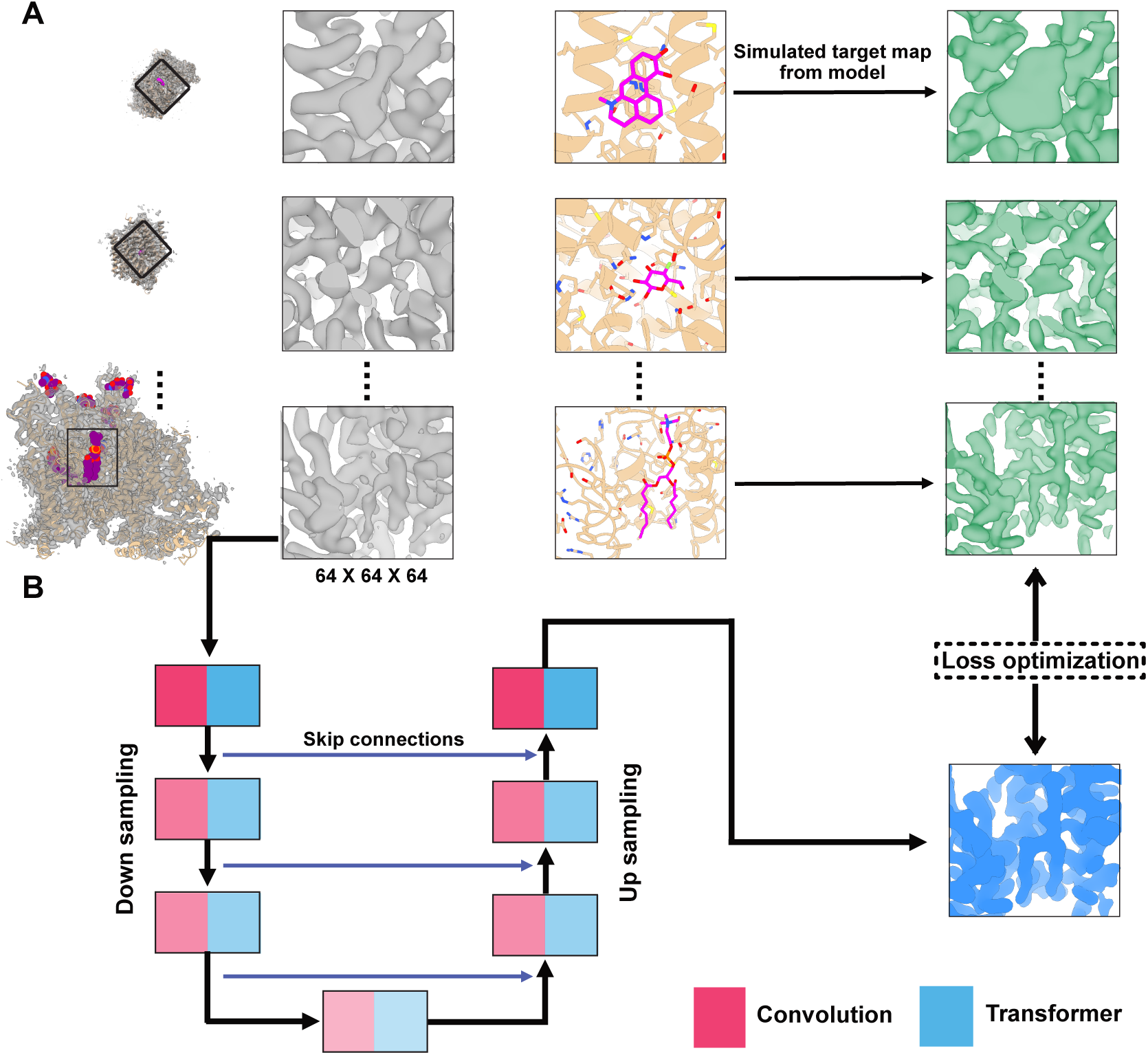
Overview of CryoLigATE. (A) Data preparation pipeline. Experimental cryo-EM maps are extracted as 64 *×* 64 *×* 64 voxel volumes centered on the ligand binding pocket. Example data points from EMDB/PDB ID shown are: 67602/21DU, 33962/7YNI, and 6996/6A91 for the top, middle and bottom, respectively. Simulated target maps (green) are generated from atomic models to serve as the ground truth for density sharpening. (B) Schematic of the CryoLigATE neural network architecture. The model utilizes a hybrid U-Net design featuring skip connections and alternating Convolution (red) and Transformer (blue) processing blocks. The model is trained via loss optimization by comparing the predicted sharpened density (blue) against the simulated target map.

### Quantitative enhancement of map resolvability by CryoLigATE

To evaluate the performance of CryoLigATE on our independent test set (n = 649), we calculated the Q-score [16], a quantitative metric that measures how clearly individual atoms or residues are resolved in a 3D density map, for each deposited atomic model and our predicted map. We compared them against the baseline Q-scores of the original deposited maps. Like the full dataset, the test dataset contains diverse ligand types including drugs, lipids, and steroids (Fig. S5). Although we did not observe significant improvement for the overall pocket (ligand + protein) and ligand-only densities (Fig. 4, *left*), this comparison could be influenced by maps in which the ligand density was already of high quality; therefore, we further investigated whether the model specifically improved targets with initially poor input densities. For a subset of data points where the deposited map had a ligand-only Q-score below 0.5 (n = 81), we found that the predicted maps showed a significant improvement (Fig. 4, *right*), demonstrating the method’s capacity for enhancement when operating on inputs of low quality. The change in ligand Q-score (ΔQ = predicted - input) exhibited a broadly positive distribution, ranging from -0.05 to 0.3 for the ligand alone, quantifying the model’s ability to enhance map quality (Fig. S6). We did not observe significant improvement in the surrounding protein residues, possibly because our loss function places greater emphasis on the ligand density map (see Methods for more details).

We further sought to determine whether the model ever degraded map quality. First, we identified poorly predicted maps by extracting data points where the ligand Q-score for the predicted maps fell below 0.5 (n = 58). Comparing these against their original deposited inputs revealed no significant change in the distribution, indicating that poor predictions were generally inherited from poor inputs rather than introduced by the model (Fig. S7, *top*). Finally, we examined targets with highly resolved initial maps (deposited ligand Q-score *>* 0.5, n = 558) and observed no significant reduction in their predicted Q-scores (Fig. S7, *top*). This confirms that CryoLigATE does not degrade high-quality inputs, preserving the integrity of well-resolved experimental densities.

**Fig. 4:**
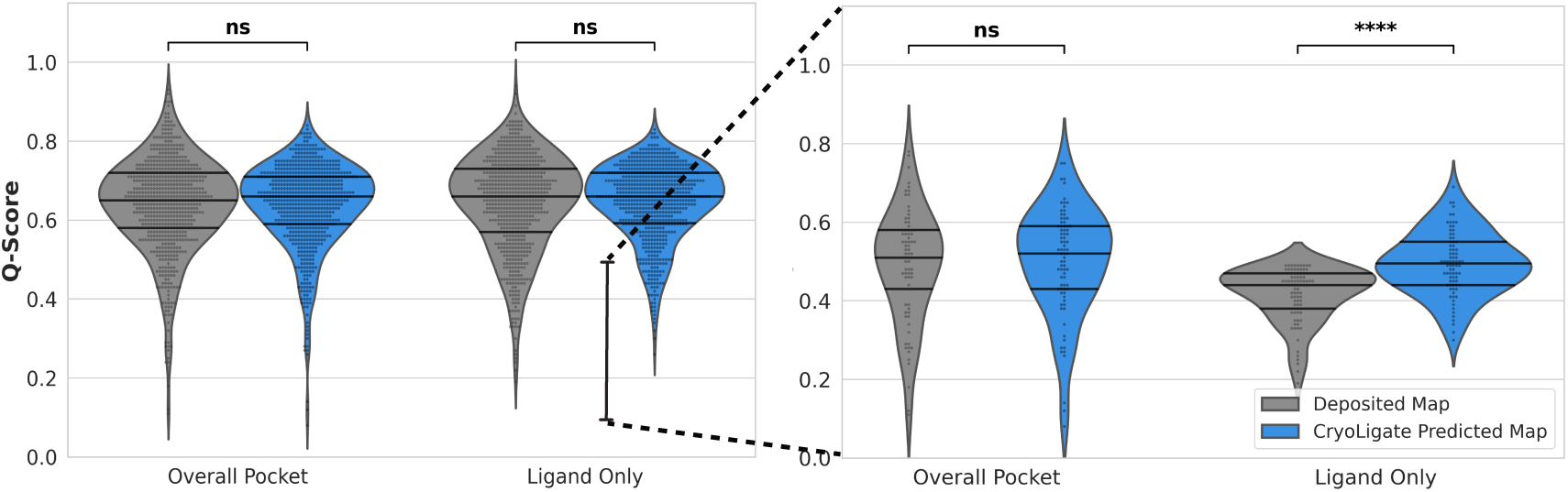
Quantitative enhancement of map resolvability by CryoLigATE. (*Left*) Comparison of model-to-map Q-scores between the original deposited experimental maps (gray) and the CryoLigATE predicted maps (blue) for the testing dataset (n = 649). Distributions are shown as violin and swarm plots for both the “Overall Pocket” (protein and ligand environment) and the “Ligand Only” density. Statistical significance is indicated by asterisks (*∗ ∗ ∗∗* denotes p*≤*0.0001). (*Right*) The same analysis focused on the poor Q-score values for the ligand (p*≤*0.5, n = 81) for the deposited model, as highlighted by the line segment on the left plot.

### CryoLigATE enhances ligand resolvability across diverse targets

To evaluate the impact of CryoLigATE on experimental map quality, we examined representative protein-ligand complexes from the independent test set. In all instances, CryoLigATE-processed maps demonstrated marked improvements in feature contrast and resolvability for the ligand (Fig. 5).

As also illustrated in Fig. 1, the short prokaryotic Argonaute protein complex MapSPARTA defends bacteria by inducing cell death in the presence of pathogenic nucleotides [14]. In a structure of the 504-kD soluble dimer bound to NAD^+^ (EMD-40680, PDB 8SPO), CryoLigATE improved the ligand *Q*-score from 0.19 to 0.41 (Fig. 5A), and resolved previously discontinuous densities into a continuous envelope.

As an example of a multimeric viral membrane protein bound to a sugar molecule, influenza hemagglutinin binds sialic acids tethered to glycoproteins on the surface of host cells, for example in the respiratory tract [17]. In a structure of the 172-kD hemagglutinin trimer from H2N2 avian influenza bound to *N* -acetyl-*α*-neuraminic acid (EMD-38603, PDB 8XRD), CryoLigATE increased the ligand *Q*-score from 0.44 to 0.65 (Fig. 5B), clarifying topology particularly around the acetamido group.

As an example of a multimeric human membrane protein bound to a lipidic modulator, the endogenous neurosteroid allopregnanolone recently became the first FDA-approved drug to treat post-partum depression, by enhancing the activity of *γ*-aminobutyric acid type A (GABA_A_) receptors [18]. In a structure of the 365-kD pentameric receptor (EMD-40503, PDB 8SI9), the allopregnanolone *Q*-score increased from 0.34 to 0.56, with clear distinguishability from neighboring protein residues (Fig. 5C).

As an illustrative complex involving a monomeric membrane protein, the sphingosine-1-phosphate receptor regulates homeostasis and various disease states [19]. Among other things, it is targeted by siponimod, a prescription medication used to treat relapsing multiple sclerosis. In a structure of the 159-kD G-protein complex (EMD-31344, PDB 7EW1), CryoLigATE improved the siponimod *Q*-score from 0.22 to 0.46 (Fig. 5D). Visually, this improvement was reflected in enhanced resolvability especially around the trifluoromethyl group.

Finally, in an example of a large lipidated multimeric mammalian membrane protein, the mitochondrial acyl carrier protein plays a critical structural role in respiratory complex I, locking together adjacent components via a 4’-phosphopantetheine tethered fatty acid [20]. In a structure of the 614-kD membrane arm of ovine respiratory complex I (EMD-4479, PDB 6Q9B), CryoLigATE increased the *Q*-score for the tethered S-tetradecanoyl-4’-phosphopantetheine from 0.38 to 0.56 (Fig. 5E), enabling topological continuity from the phosphate linker through the hydrophobic tail.

Collectively, these results illustrate various improvements by CryoLigATE in quantitative and visual resolvability of volumetric ligand representations in experimental cryo-EM maps. The enhanced interpretability of structural information across diverse ligand classes, including small drug-like molecules, sugars, and lipids, highlights the method’s utility in generating high-confidence maps for precise atomic model building and the study of molecular recognition.

**Fig. 5:**
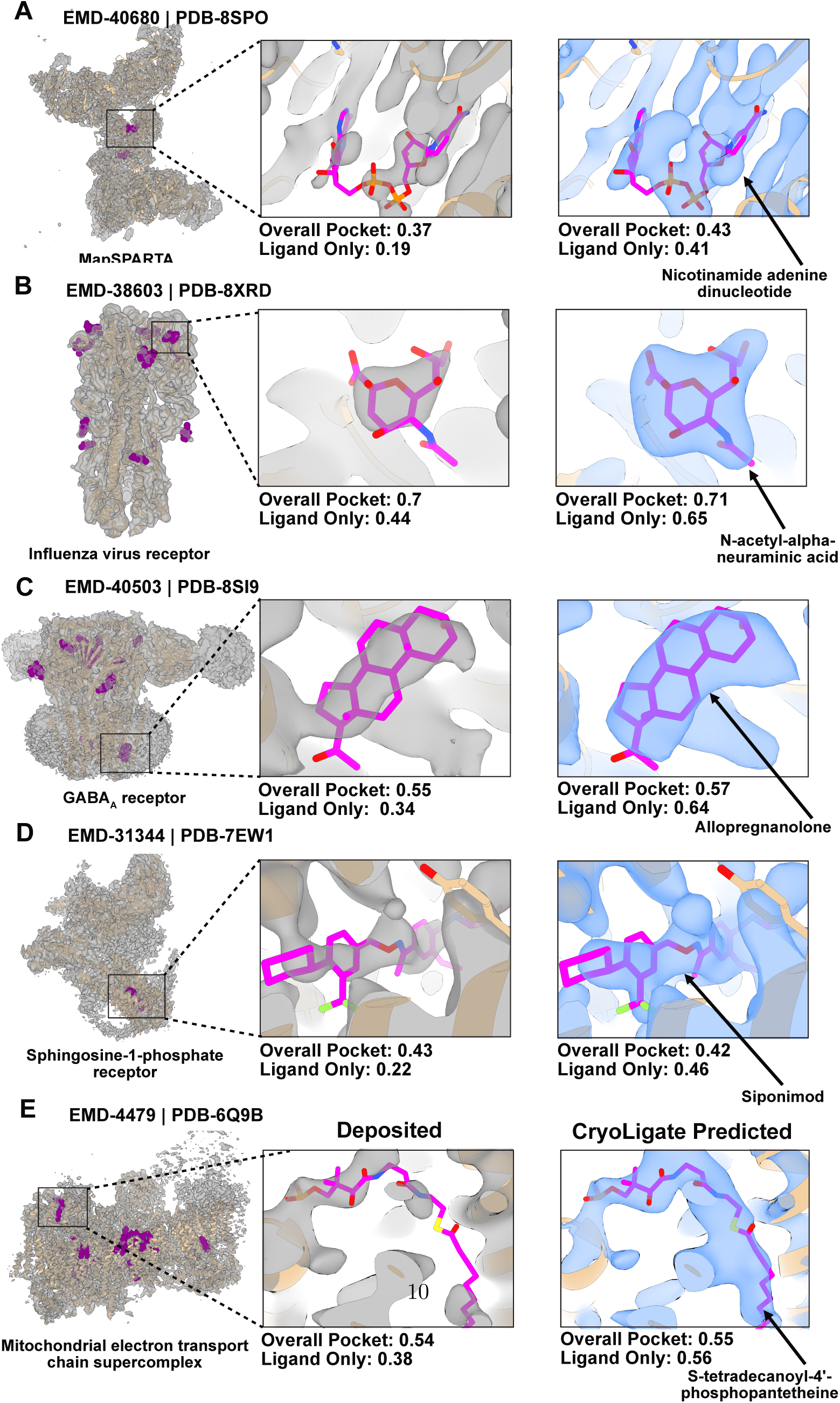
CryoLigATE enhances ligand resolvability across diverse targets. (A-E) Visual and quantitative comparison between deposited experimental cryo-EM maps and CryoLigATE-predicted maps across five representative complexes. *Left* Global view of the experimental density (gray) and modeled protein (tan ribbons) with modeled ligands highlighted (magenta spheres). *Center* Magnified view of the ligand region in the “Deposited” experimental density (gray) subjected to CryoLigATE processing. *Right* Magnified view of the “CryoLigATE Predicted” density (blue) after processing. Quantification via Q-scores (Overall Pocket and Ligand Only) is indicated below each inset. In magnified views, the target ligand and proximal side chains are shown as sticks, colored by heteroatom.

### Robustness of the approach

During CryoLigATE inference, although the user provides an aligned PDB model containing the ligand, this coordinate information is utilized solely to localize the region of interest. Only the localized experimental density and the derived protein occupancy mask are used as input for CryoLigate. To assess the robustness and algorithmic stability of our method, we subjected CryoLigATE to a series of coordinate perturbation tests. Using the well-defined density of the GABA*_A_* receptor bound to allopregnanolone (EMD-40503) as a baseline control, we deliberately introduced spatial and geometric noise to the input ligand prior to map processing. Specifically, the initial ligand models were subjected to a rotation of 90°, translational shift of 3 Å, and substitution of the ligand by a single carbon atom prior to map processing (Fig. S8A,B,C). Across these perturbed conditions, CryoLigATE demonstrated consistent recovery of the expected ligand density, despite the initial coordinate inaccuracies. To investigate limitations of the algorithm, particularly the risk of artificial density generation, we also performed a negative control experiment by directing CryoLigATE to process a volumetric region devoid of experimental density (Fig. S8D). Under these conditions, CryoLigATE did produce an unstructured, amorphous density centered in the target region, indicating a susceptibility to map hallucination in the absence of input signal. This result underscores a critical caveat: CryoLigATE is designed strictly to refine pre-existing, putative ligand densities, and should not be used to probe or generate density in visibly empty pockets.

## Discussion

In this study, we demonstrate that deep learning can be used to significantly improve feature contrast and interpretability for small-molecule ligands within experimental cryo-EM density maps where initial ligand map quality is poor. The method was trained and validated across a chemically diverse landscape, including pharmaceuticals, lipids, carbohydrates, and steroids, spanning a broad range of molecular weights, lipophilicities, and conformational degrees of freedom (Figs. S1 and 2). Our results highlight a systematic enhancement of previously ill-defined or discontinuous density features, yielding continuous envelopes for flexible ligand regions and clearly resolved chemical functional groups. By clarifying these localized volumes, the method provides a more robust empirical baseline for building confident structural models.

It is important to contextualize our approach within the rapidly evolving landscape of structural bioinformatics. Current state-of-the-art models, such as AlphaFold3 [21] and Boltz-2 [22], have made extraordinary advances by predicting protein-ligand complexes from amino acid sequences and ligand SMILES strings alone. However, when predicting novel interactions without direct experimental constraints, these models can face challenges, including the occasional hallucination of binding modes or deviations from expected physical behaviors, such as steric clashes or an inability to accurately respond to disruptive protein mutations [23, 24]. The distinct advantage of CryoLigATE is that it avoids blind prediction by anchoring deep spatial and structural priors directly to experimental data. Our model provides a data-driven refinement of experimental density rather than a purely statistical inference.

Crucially, CryoLigATE does not rely on user-provided SMILES strings or chemical identifiers during the inference process. By operating independently of external chemical input, the method minimizes the risk of user-induced bias and further reduces the potential for structural hallucinations. This map-centric approach ensures that the resulting density is a true reflection of the experimental observation, refined by the network’s learned understanding of molecular physics, geometry, and general properties of ligands bound to macromolecular structures.

A limitation of the current CryoLigATE method is the treatment of the ligand as a single, static entity. In biological systems, ligand binding is often heterogeneous, characterized by conformational flexibility and variable fractional occupancy. Because CryoLigATE operates on processed consensus cryo-EM maps and deposited atomic coordinates, it is designed to predict the best representative state of the densities captured within that reconstruction. Modeling the continuous distribution of ligand poses or identifying discrete, low-occupancy sub-states remains a major challenge. Addressing this in the future may require algorithms that bypass consensus maps entirely to extract structural heterogeneity directly from raw single-particle images or localized micrograph regions [25–27].

Furthermore, our investigations highlight a fundamental bottleneck in training AI models for cryo-EM: the volume of high-resolution protein-ligand complexes is currently limited, and deposited models may contain coordinate errors if originally built into lower-resolution densities. To tackle these limitations, a data-augmentation approach leveraging high-confidence complexes from X-ray crystallography databases, such as PDBBind [28], could provide high-fidelity ground truths. However, generating realistic low-resolution inputs that accurately recapitulate the inherent noise and artifacts of cryo-EM reconstruction remains a challenge. This suggests a need for novel methodological developments in physics-based simulations or generative approaches, where models like cryoGAN have shown promise [29].

Looking forward, the principles underlying CryoLigATE hold promise for other modalities, such as cryo-electron tomography (cryo-ET) and time-resolved cryo-EM, where signal-to-noise ratios are notoriously low, potentially accelerating structure-based drug discovery and our understanding of dynamic biological processes *in situ*. As deep learning tools for map enhancement become standard practice, the structural biology community may benefit from establishing clear guidelines for their application. We suggest that AI-enhanced maps serve best as complementary interpretive aids to guide manual model building, confirm binding pockets, and facilitate initial ligand refinement. To ensure long-term data integrity, final atomic refinement should ideally remain tethered to the original experimental densities. Furthermore, we encourage the practice of depositing both the original and the post-processed maps in the EMDB. Because future AI models will inevitably learn from deposited data, maintaining access to the raw experimental signal is vital to prevent model collapse - a scenario where recursive training on AI-generated outputs could inadvertently amplify structural biases. By grounding to the primary experimental data, the field can ensure that AI continues to serve as a tool for illuminating, rather than obscuring the underlying biological truths.

## Methods

### Data curation

Training neural networks to enhance 3D density maps requires a large, well-curated dataset of paired low- and high-resolution protein–ligand complexes. The former serve as inputs and the latter as ground-truth targets for the AI framework. We treated raw, experimental cryo-EM maps as “low-resolution” inputs, and used forward-simulated 3D EM maps derived from their deposited atomic models as the “high-resolution” targets, a strategy previously successful in denoising protein maps [10].

Initial structure metadata was sourced from the Research Collaboratory for Structural Bioinformatics (RCSB) Protein Data Bank (PDB) using a combination of Search, REST, and GraphQL APIs [30]. The dataset was restricted to structures determined by electron microscopy with a reported global resolution of 4.0 Åor better, deposited on or before March 31st, 2026. Common monoatomic and polyatomic ions (e.g., Zn^2+^, Mg^2+^, SO^2^*^−^*), crystallization buffers (e.g., HEPES, TRIS), and detergents (e.g., DDM, BOG) were removed.

To prevent model over-fitting and mitigate the overrepresentation of common targets, we applied a two-stage deduplication protocol. First, we retained only unique protein-ligand pairs per PDB entry, defined by unique combinations of PDB ID and ligand CCD code. Second, we addressed chronological and target-centric bias by capping the dataset at a maximum of 50 structurally diverse representative samples per unique ligand species. This ensured the AI framework remained robust across a broad chemical space rather than specializing in high-frequency scaffolds. Following metadata retrieval, the corresponding atomic coordinate files (.cif) and their associated un-masked 3D volumetric maps (.map.gz) were automatically downloaded from the RCSB PDB [30] and the Electron Microscopy Data Bank (EMDB) via the unified wwPDB file servers.

### Cryo-EM Map Processing

Raw cryo-EM maps and their corresponding atomic models were processed to extract localized, protein-ligand structural environments. For each target ligand, a localized bounding region centered on the ligand’s Cartesian centroid was isolated from the native experimental density map. To ensure absolute spatial fidelity and prevent geometric distortions across heterogeneous datasets, target grid points were defined in absolute physical space (Å) and mapped back to the native density coordinates using the fractional transformation matrix (**M**_frac_) derived from the map’s unit cell parameters.

This isolated physical sub-volume was then computationally resampled to a uniform isotropic voxel size of 0.5 Å using third-order cubic spline interpolation via the Gemmi library [31], yielding a standardized 3D cubic grid of 64 *×* 64 *×* 64 voxels (32^3^ Å^3^). This workflow mathematically preserves uniform physical dimensions (e.g., secondary structure widths and bond distances) across all structural volumes, irrespective of the native experimental resolution.

To eliminate spatial ambiguity during model training, a strict spatial filter was applied; grid environments containing more than one distinct ligand density were discarded. Additionally, ligands with a maximum atomic diameter exceeding the bounding box dimensions (minus a conservative 4.0 Å spatial buffer) were excluded to prevent boundary clipping effects. This filtering workflow yielded a final dataset of 6,511 unique protein-ligand complex cryo-EM densities.

### Synthetic Ground Truth Generation

To establish high resolution “ground truth” targets for the AI model to optimize against, synthetic density maps were computationally generated for each complex. The deposited atomic coordinates of the protein and ligand were projected onto the 64 × 64 × 64 voxel grid using a 3D Gaussian filter. We utilized a standard cryo-EM distribution factor (*σ ≈* 0.225 *×* resolution). Finally, both the experimental cryo-EM sub-volumes and the synthetic ground truth maps were subjected to strictly paired *Z*-score normalization (zero mean and unit variance) before being serialized into high-performance HDF5 format for distributed deep learning.

### Neural Network Architecture

To perform density enhancement, we utilized a customized 3D Swin-Conv UNet (SCUNet) architecture [15]. This model adopts a hierarchical U-Net topology consisting of three downsampling and upsampling stages, integrated via additive skip connections to preserve multi-scale structural information. The network accepts a single-channel 3D input tensor (experimental density) and outputs a single-channel enhanced density map.

The core of the architecture is the ConvTransBlock, which employs a “split-and-aggregate” strategy to process features. Upon entering the block, the input feature map is split along the channel dimension into two symmetric paths. A local feature extractor consisting of two 3 *×* 3 *×* 3 convolutions, each followed by Filter Response Normalization (FRN) and a Thresholded Linear Unit (TLU). This path captures high-frequency local textures and enforces translational equivariance. A global contextual branch utilizing 3D Windowed Multi-head Self-Attention (WMSA). This path models long-range dependencies by computing self-attention within 4 *×* 4 *×* 4 local windows. To enable cross-window communication, we alternated between standard Windowed Attention (W-MSA) and Shifted Window Attention (SW-MSA). In shifted layers, the window partitions are displaced by half the window size (2 *×* 2 *×* 2 voxels), with a masking mechanism applied to handle boundary conditions. To incorporate spatial geometry, we utilized a 3D Relative Position Bias, where a learnable parameter matrix is indexed by the relative coordinate distance between voxels within each window, significantly improving the network’s ability to learn consistent protein-ligand geometries.

To maintain robust performance across varied cryo-EM density scales and small batch sizes, we replaced standard Batch Normalization in the convolutional blocks with Filter Response Normalization (FRN). Unlike batch-dependent methods, FRN normalizes across spatial dimensions independently for each channel (*H × W × D*), using a learnable threshold *τ* to prevent vanishing activations.

### Loss function

Standard pixel-wise loss functions (e.g., Mean Squared Error) often cause deep learning models to output highly conservative, low-intensity density maps. Such outputs inherently score poorly on structural biology metrics like the Q-score [16]. We utilized a Loss (*L_T otal_*) formulated as a weighted sum of three penalty terms:

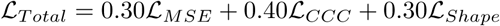

Spatially-Weighted Mean Squared Error (*L_MSE_*): To anchor global density while forcing the network to prioritize the sparsely populated ligand pocket, we applied a dynamic spatial weight map (*W*). Voxels residing within the ground-truth ligand mask (*M*) were penalized with a 30-fold multiplier compared to the protein and solvent background:

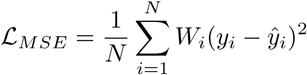

where *W_i_* = 30 if *M_i_ >* 0.5, and *W_i_* = 1 otherwise.

Concordance Correlation Coefficient (*L_CCC_*): To ensure the predicted ligand density maintains the precise physical scale and variance of the target, we applied the Concordance Correlation Coefficient (CCC) [**?**] strictly to the ligand region (*M >* 0.5). Unlike standard Pearson correlation, CCC penalizes deviations in both mean intensity and amplitude variance:

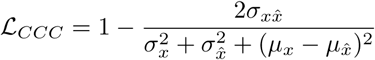

Global Dice and Ligand Tversky Loss (*L_Shape_*): To ensure 3D boundary overlap, we utilized a composite shape loss:

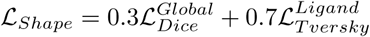

The Tversky loss [32] for the ligand used parameters *α* = 0.2 and *β* = 0.8. This heavy skew penalizes false negatives, forcing the network to favor continuous, unbroken density over sparse, disjointed predictions.

### Training and Evaluation pipeline

Training Pipeline and EvaluationThe dataset was randomly shuffled using a fixed seed to eliminate chronological and detector-based hardware biases, ensuring that older, lower-resolution structures were evenly distributed across the 80/10/10 (Train/Validation/Test) splits. To establish independent training, validation, and testing partitions free from homology-induced bias, we implemented a ligand-specific, sequence-based clustering workflow. For each unique ligand class, an exact N-by-N pairwise comparison was performed among all associated PDB structures. Global sequence alignments were computed across all extracted polymer chains, and the structural similarity between any two PDB entries was defined as the maximum sequence identity observed between any of their respective chain pairs. Using a graph-theoretic approach, an edge was mapped between PDB entries sharing a maximum chain sequence identity of 40% or higher, and discrete homology groups were resolved by extracting the graph’s connected components. Structures without homologous partners or with missing sequence data were designated as isolated singletons. By enforcing that all members of a given sequence cluster are strictly assigned to the same partition, we guarantee that the train, validation, and test sets are mutually exclusive, preventing data leakage and ensuring a rigorous evaluation of the model.

The model was trained for 200 epochs on NVIDIA A100 GPUs using the AdamW optimizer [33] with an initial learning rate of 1 *×* 10*^−^*^4^ and a weight decay of 1 *×* 10*^−^*^2^. We employed a ReduceL-ROnPlateau scheduler to halve the learning rate upon validation stagnation. To improve model generalization, we implemented stochastic 3D data augmentation, including random axial flips and 90-degree rotations along the (XY, XZ, YZ) planes. Because raw loss values do not always correlate with biological map quality, during training and validation, a sigmoid activation was applied to the output; however, during final inference, this squashing was omitted. The raw logits were directly mapped to the original physical unit cell and orthogonal origins using the Gemmi library. This preserves the physical dynamic range of the density, ensuring that the resulting .mrc files are fully compatible with downstream validation tools and visualization in ChimeraX [34].

### Inference Pipeline

The inference pipeline is designed as a standalone command-line tool, allowing for the targeted enhancement of any ligand within a cryo-EM map. It takes about a second to process one complex in a NVIDIA GenForce RTX 3070 GPU desktop. The syntax for executing the refinement is as follows:

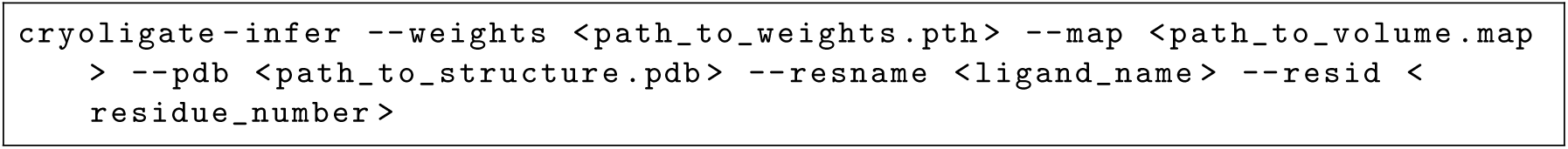

Although the user provides an aligned PDB model containing the ligand, this coordinate information is utilized solely to localize the region of interest. Only the localized experimental density and the derived protein occupancy mask are used as input for CryoLigATE.

## Acknowledgments

We acknowledge Elisei Mankov, Archit Kumar Vasan, and Piotr Draczkowski for discussion and suggestions in the project. CryoLigATE training was performed using the Berzelius supercomputing resources which is a part of National Academic Infrastructure for Supercomputing in Sweden (Berzelius-2026-98; NAISS 2025/3-71). This work was partially funded through a Marie Sklodowska-Curie Postdoctoral Fellowship 101107036 to NH, and grants from the Swedish Research Council (VR; 2019-02433, 2025-06231), the Knut and Alice Wallenberg foundation (KAW; 2023.0254), the BioExcel-3 Centre-of-Excellence (EuroHPC Joint Undertaking; 101093290), and the Swedish e-Science Research Center to EL.

## Data and code availability

The source code for model inference along with the pretrained neural network weights are hosted on GitHub (https://github.com/nandanhaloi123/CryoLigATE). The complete data pipeline, including the class-balanced 3D volumetric HDF5 training dataset, statistical inventories, and model training code, have been deposited repository-wide to Zenodo (doi:10.5281/zenodo.20794212).

## Author Contributions

**Conceptualization: NH**

**Data curation: NH**

**Formal analysis: NH**

**Funding acquisition: NH, EL**

**Investigation: NH**

**Methodology: NH**

**Project administration: RJH, EL**

**Resources:NH, EL**

**Software:NH**

**Supervision:RJH, EL**

**Validation: NH**

**Visualization: NH**

**Writing – original draft: NH**

**Writing – review & editing: NH, RJH, EL**

**Declaration of interests. The authors declare no competing interests.**

## References

[1] Bertram, K., Agafonov, D.E., Dybkov, O., Haselbach, D., Leelaram, M.N., Will, C.L., Urlaub, H., Kastner, B., Lührmann, R., Stark, H.: Cryo-EM structure of a pre-catalytic human spliceosome primed for activation. Cell 170(4), 701–713 (2017)

[2] Amunts, A., Brown, A., Toots, J., Scheres, S.H., Ramakrishnan, V.: The structure of the human mitochondrial ribosome. Science 348(6230), 95–98 (2015)

[3] Kim, J.J., Gharpure, A., Teng, J., Zhuang, Y., Howard, R.J., Zhu, S., Noviello, C.M., Walsh Jr, R.M., Lindahl, E., Hibbs, R.E.: Shared structural mechanisms of general anaesthetics and benzodiazepines. Nature 585(7824), 303–308 (2020)

[4] Bartesaghi, A., Aguerrebere, C., Falconieri, V., Banerjee, S., Earl, L.A., Zhu, X., Grigorieff, N., Milne, J.L., Sapiro, G., Wu, X., et al.: Atomic resolution cryo-EM structure of *β*-galactosidase. Structure 26(6), 848–856 (2018)

[5] Haloi, N., Howard, R.J., Lindahl, E.: Cryo-em ligand building using alphafold3-like model and molecular dynamics. PLOS Computational Biology 21(8), 1013367 (2025)

[6] Muenks, A., Zepeda, S., Zhou, G., Veesler, D., DiMaio, F.: Automatic and accurate ligand structure determination guided by cryo-electron microscopy maps. Nature Communications 14(1), 1164 (2023)

[7] Sweeney, A., Mulvaney, T., Maiorca, M., Topf, M.: ChemEM: flexible docking of small molecules in cryo-em structures. Journal of medicinal chemistry 67(1), 199–212 (2023)

[8] Robertson, M.J., Zundert, G.C., Borrelli, K., Skiniotis, G.: GemSpot: a pipeline for robust modeling of ligands into cryo-EM maps. Structure 28(6), 707–716 (2020)

[9] Vant, J.W., Lahey, S.-L.J., Jana, K., Shekhar, M., Sarkar, D., Munk, B.H., Kleinekathöfer, U., Mittal, S., Rowley, C., Singharoy, A.: Flexible fitting of small molecules into electron microscopy maps using molecular dynamics simulations with neural network potentials. Journal of chemical information and modeling 60(5), 2591–2604 (2020)

[10] He, J., Li, T., Huang, S.-Y.: Improvement of cryo-em maps by simultaneous local and non-local deep learning. Nature Communications 14(1), 3217 (2023)

[11] Wang, X., Zhu, H., Terashi, G., Taluja, M., Kihara, D.: Diffmodeler: large macromolecular structure modeling for cryo-em maps using a diffusion model. Nature methods 21(12), 2307– 2317 (2024)

[12] Sanchez-Garcia, R., Gomez-Blanco, J., Cuervo, A., Carazo, J.M., Sorzano, C.O.S., Vargas, J.: DeepEMhancer: a deep learning solution for cryo-em volume post-processing. Communications biology 4(1), 874 (2021)

[13] Berkeley, R.F., Cook, B.D., Herzik Jr, M.A.: Machine learning approaches to cryo-em density modification differentially affect biomacromolecule and ligand density quality. Frontiers in Molecular Biosciences 11, 1404885 (2024)

[14] Shen, Z., Yang, X.-Y., Xia, S., Huang, W., Taylor, D.J., Nakanishi, K., Fu, T.-M.: Oligomerization-mediated activation of a short prokaryotic argonaute. Nature 621(7977), 154–161 (2023)

[15] Liu, Z., Lin, Y., Cao, Y., Hu, H., Wei, Y., Zhang, Z., Lin, S., Guo, B.: Swin transformer: Hierarchical vision transformer using shifted windows. In: Proceedings of the IEEE/CVF International Conference on Computer Vision, pp. 10012–10022 (2021)

[16] Pintilie, G., Zhang, K., Su, Z., Li, S., Schmid, M.F., Chiu, W.: Measurement of atom resolvability in cryo-em maps with Q-scores. Nature methods 17(3), 328–334 (2020)

[17] Sun, J., Zheng, T., Jia, M., Wang, Y., Yang, J., Liu, Y., Yang, P., Xie, Y., Sun, H., Tong, Q., et al.: Dual receptor-binding, infectivity, and transmissibility of an emerging H2N2 low pathogenicity avian influenza virus. Nature Communications 15(1), 10012 (2024)

[18] Legesse, D.H., Fan, C., Teng, J., Zhuang, Y., Howard, R.J., Noviello, C.M., Lindahl, E., Hibbs, R.E.: Structural insights into opposing actions of neurosteroids on GABAA receptors. Nature Communications 14(1), 5091 (2023)

[19] Yuan, Y., Jia, G., Wu, C., Wang, W., Cheng, L., Li, Q., Li, Z., Luo, K., Yang, S., Yan, W., et al.: Structures of signaling complexes of lipid receptors S1PR1 and S1PR5 reveal mechanisms of activation and drug recognition. Cell research 31(12), 1263–1274 (2021)

[20] Letts, J.A., Fiedorczuk, K., Degliesposti, G., Skehel, M., Sazanov, L.A.: Structures of respiratory supercomplex I+ III2 reveal functional and conformational crosstalk. Molecular cell 75(6), 1131–1146 (2019)

[21] Abramson, J., Adler, J., Dunger, J., Evans, R., Green, T., Pritzel, A., Ronneberger, O., Willmore, L., Ballard, A.J., Bambrick, J., et al.: Accurate structure prediction of biomolecular interactions with alphafold 3. Nature, 1–3 (2024)

22. Passaro, S., Corso, G., Wohlwend, J., Reveiz, M., Thaler, S., Somnath, V.R., Getz, N., Portnoi, T., Roy, J., Stark, H., et al.: Boltz-2: Towards accurate and efficient binding affinity prediction. BioRxiv (2025)

[23] Masters, M.R., Mahmoud, A.H., Lill, M.A.: Investigating whether deep learning models for co-folding learn the physics of protein-ligand interactions. Nature Communications 16(1), 8854 (2025)

24. Buttenschoen, M., Morris, G.M., Deane, C.M.: PoseBusters: AI-based docking methods fail to generate physically valid poses or generalise to novel sequences. Chemical Science 15(9), 3130–3139 (2024)

[25] Zhong, E.D., Bepler, T., Berger, B., Davis, J.H.: CryoDRGN: reconstruction of heterogeneous cryo-em structures using neural networks. Nature methods 18(2), 176–185 (2021)

[26] Punjani, A., Fleet, D.J.: 3D variability analysis: Resolving continuous flexibility and discrete heterogeneity from single particle cryo-em. Journal of structural biology 213(2), 107702 (2021)

[27] Toader, B., Sigworth, F.J., Lederman, R.R.: Methods for cryo-em single particle reconstruction of macromolecules having continuous heterogeneity. Journal of molecular biology 435(9), 168020 (2023)

[28] Liu, Z., Li, Y., Han, L., Li, J., Liu, J., Zhao, Z., Nie, W., Liu, Y., Wang, R.: PDB-wide collection of binding data: current status of the pdbbind database. Bioinformatics 31(3), 405–412 (2015)

29. Gupta, H., McCann, M.T., Donati, L., Unser, M.: CryoGAN: A new reconstruction paradigm for single-particle cryo-em via deep adversarial learning. IEEE Transactions on Computational Imaging 7, 759–774 (2021)

[30] Burley, S.K., Bhikadiya, C., Bi, C., Bittrich, S., Chen, L., Crichlow, G.V., Christie, C.H., Dalenberg, K., Di Costanzo, L., Duarte, J.M., et al.: Rcsb protein data bank: powerful new tools for exploring 3d structures of biological macromolecules for basic and applied research and education in fundamental biology, biomedicine, biotechnology, bioengineering and energy sciences. Nucleic acids research 49(D1), 437–451 (2021)

31. Wojdyr, M.: GEMMI: A library for structural biology. Journal of Open Source Software 7(73), 4200 (2022)

[32] Salehi, S.S.M., Erdogmus, D., Gholipour, A.: Tversky loss function for image segmentation using 3d fully convolutional deep networks. In: International Workshop on Machine Learning in Medical Imaging, pp. 379–387 (2017). Springer

[33] Loshchilov, I., Hutter, F.: Decoupled weight decay regularization. arXiv preprint arXiv:1711.05101 (2017)

34. Pettersen, E.F., Goddard, T.D., Huang, C.C., Meng, E.C., Couch, G.S., Croll, T.I., Morris, J.H., Ferrin, T.E.: UCSF ChimeraX: Structure visualization for researchers, educators, and developers. Protein Science 30(1), 70–82 (2021)

