## Supplementary Information for "CryoLigATE: enhancing the resolvability of cryo-EM maps in protein-ligand complexes using deep learning"

[1] SciLifeLab, Department of Biochemistry and Biophysics,
Stockholm University, Tomtebodavägen 23,
17165 Solna, Stockholm, Sweden

[2] SciLifeLab, Department of Applied Physics,
KTH Royal Institute of Technology, Tomtebodavägen 23,
17165 Solna, Stockholm, Sweden

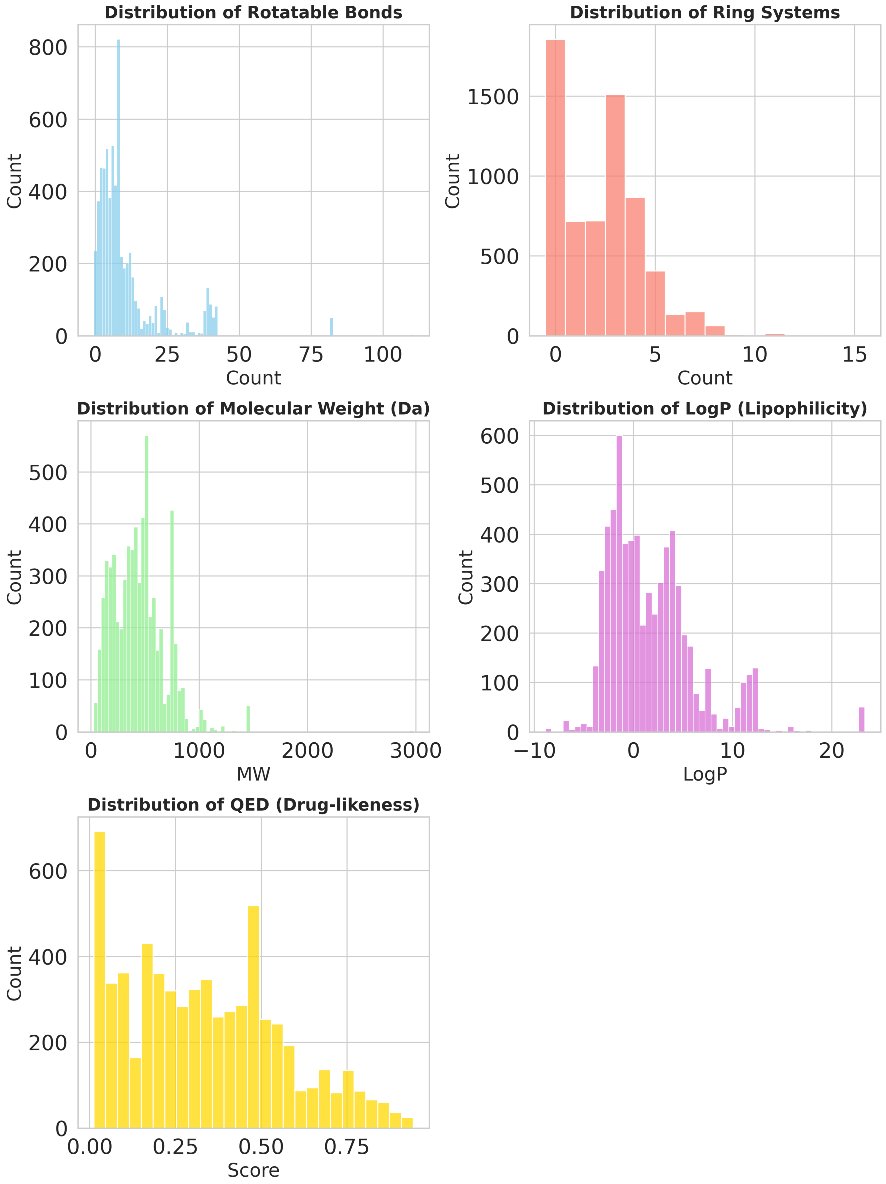


**Fig. S1**: Distribution of key physicochemical properties of the lignads, including rotatable bonds, ring systems, molecular weight, LogP, and QED scores of our curated cryo-EM protein-ligand complex dataset.

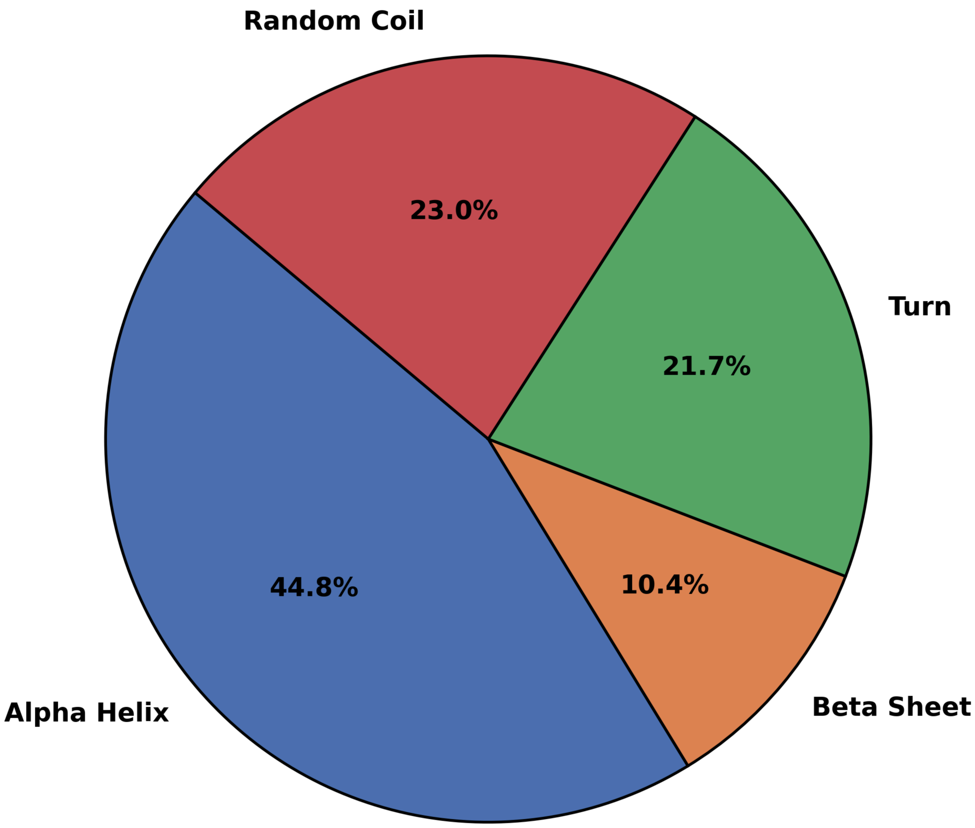


**Fig. S2**: Binding pocket secondary structure compositions.

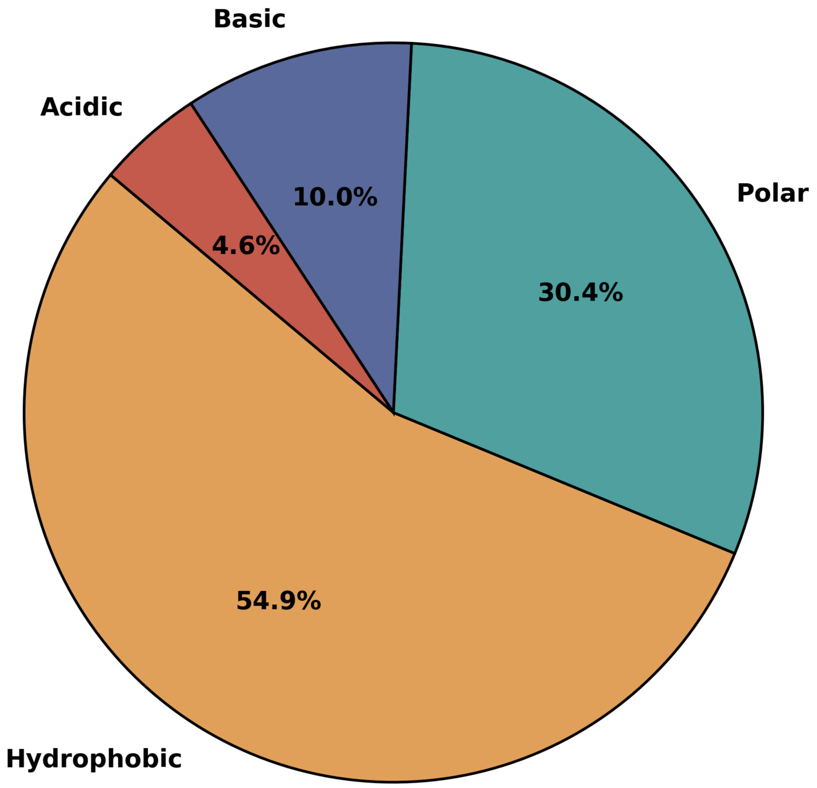


**Fig. S3**: Binding pocket chemical property compositions.

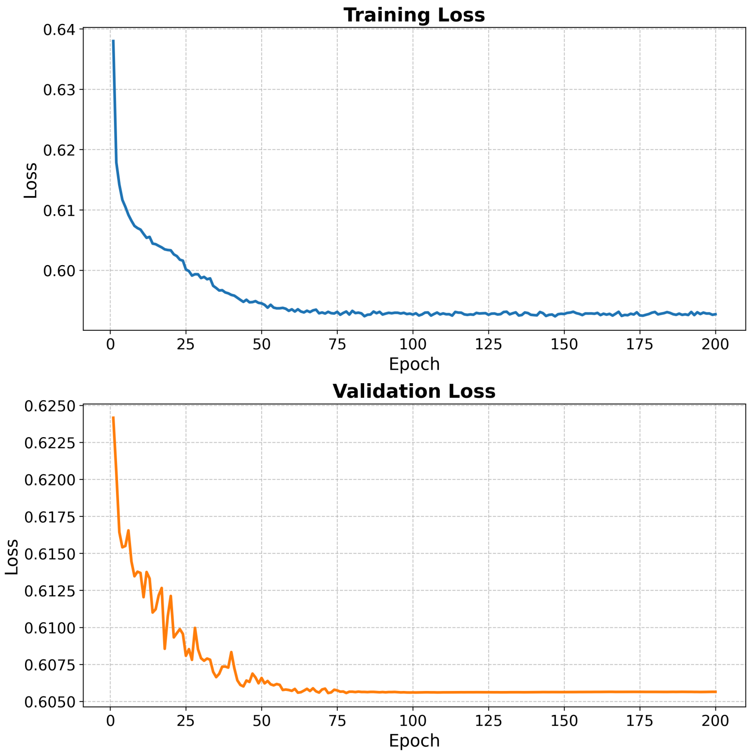


**Fig. S4**: Tracing of loss function during our training.

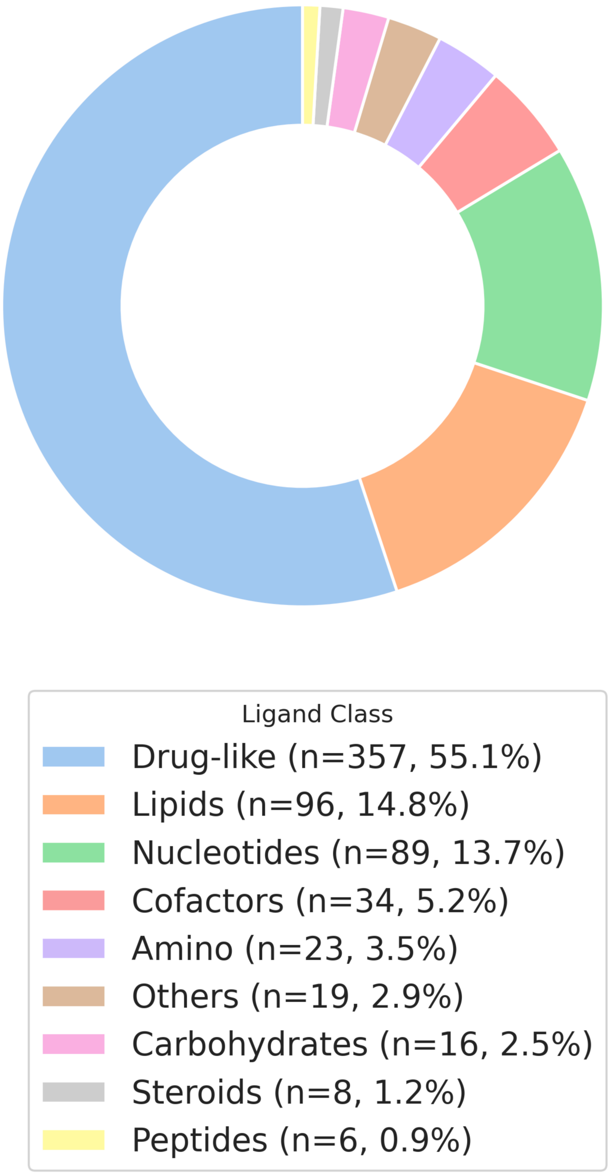


**Fig. S5**: Ligand types present in our testing dataset ($n = 648$).

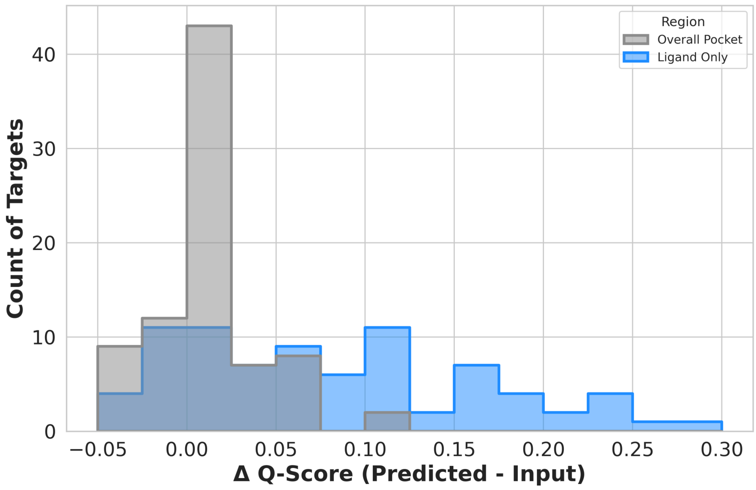


**Fig. S6**: The change in Q-score (ΔQ = predicted - input) in the testing data points where the deposited map had a ligand Q-score below 0.5 ($n = 81$).

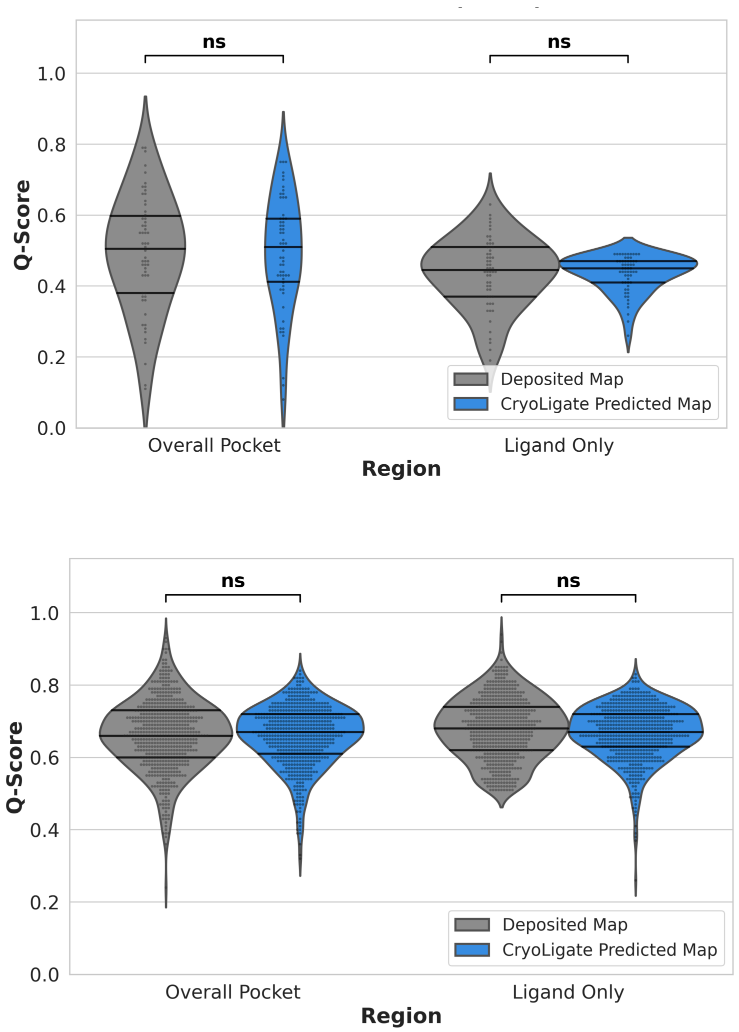


**Fig. S7**: (*Top*) Comparison of ligand Q-scores between deposited experimental maps and CryoLigATE-predicted maps for cases where the predicted ligand density exhibited low resolvability (predicted ligand Q-score $\leq 0.5$, $n = 58$). (*Bottom*) Analysis restricted to structures with highly resolved deposited ligand densities (deposited ligand Q-score $> 0.5$, $n = 558$).

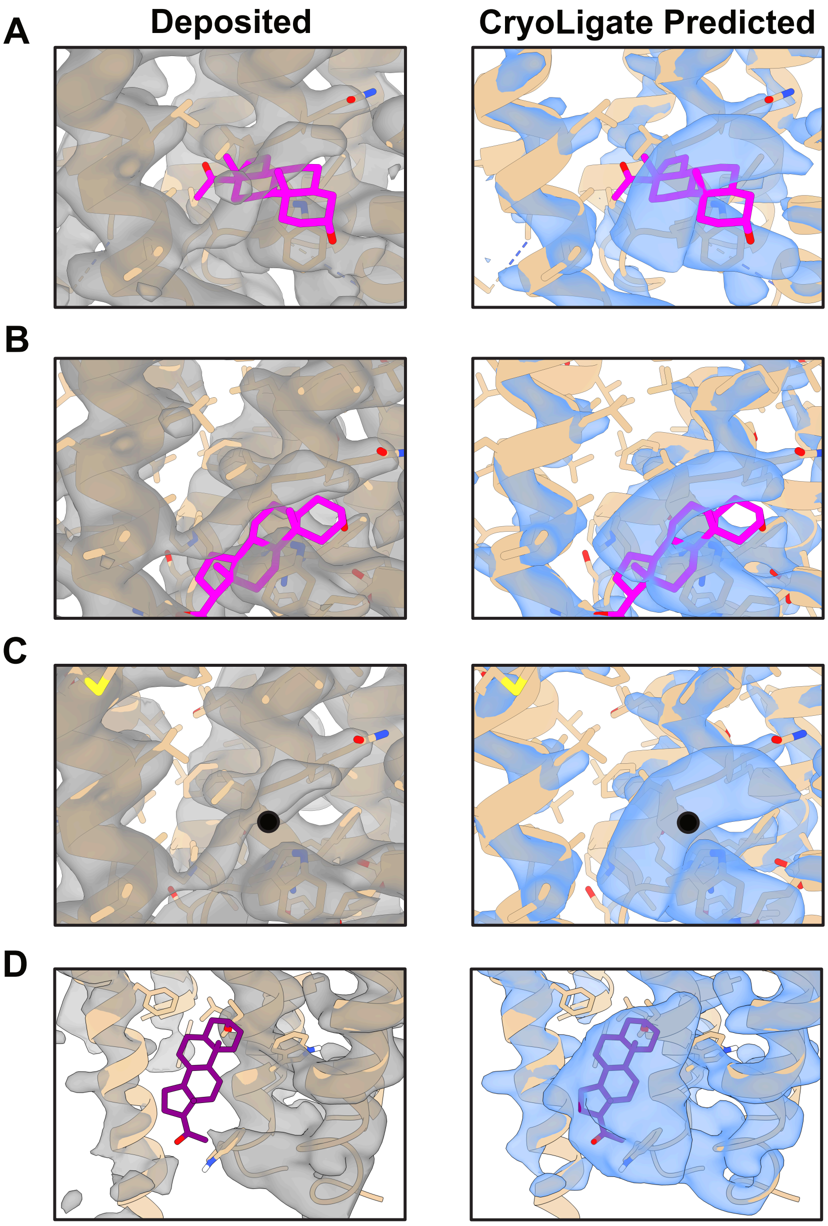


**Fig. S8**: Robustness of CryoLigATE. Algorithmic stability was evaluated using the GABA$_A$ receptor bound to allopregnanolone (EMD-40503, PDB 8SI9 [18]), a complex with well-defined experimental ligand density. To assess recovery performance, the input ligand model was deliberately subjected to (A) a rotation of 90°, (B) translational shifts of 3 Å, and (C) substitution of the ligand by a single carbon atom prior to map processing. (D) Assessment of potential map hallucination, demonstrating the algorithmic output of blobby densities when CryoLigATE is applied to a targeted region entirely lacking (Q-score ≈ 0) underlying experimental ligand signal.
